# Systematic evaluation of *de novo* mutation calling tools using whole genome sequencing data

**DOI:** 10.1101/2024.08.28.610208

**Authors:** Anushi Shah, Steven Monger, Michael Troup, Eddie KK Ip, Eleni Giannoulatou

## Abstract

*De novo* mutations (DNMs) are genetic alterations that occur for the first time in an offspring. DNMs have been found to be a significant cause of severe developmental disorders. With the widespread use of next-generation sequencing (NGS) technologies, accurate detection of DNMs is crucial. Several bioinformatics tools have been developed to call DNMs from NGS data, but no study to date has systematically compared these tools. We used both real whole genome sequencing (WGS) data from a trio from the 1000 Genomes Project (1000G) and an in-house simulated trio dataset to evaluate five DNM calling tools: DeNovoGear, TrioDeNovo, PhaseByTransmission, VarScan2, and DeNovoCNN. For DNMs called in the real dataset, we observed 8.4% concordance of variants between all tools, while 83.8% of DNMs variants were identified by only one caller. For simulated trio WGS dataset spiked with 100 DNMs, the concordance rate was also low at 3.9%. DeNovoGear achieved the highest F1 score on the real 1000G dataset, while DeNovoCNN had the highest F1 score on the simulated data. Our study provides valuable recommendations for the selection and application of DNM callers on WGS trio data.

## INTRODUCTION

*De novo* mutations (DNMs) are new mutations that arise in an individual, as opposed to being inherited from the parents. The human germline *de novo* single nucleotide variant (SNV) mutation rate is estimated at 1.0 to 1.8 x 10^-8^ which equates to 44 to 82 DNMs per genome per generation^1,2^. DNMs can occur during the formation of the gametes or post-zygotically (somatic mutations). These mutations have been associated with congenital malformations^3–5^, early-onset diseases such as intellectual disability^6^ and other developmental diseases^3,7^, and late-onset diseases such as Parkinson’s disease^8^, schizophrenia^9^, and bipolar disorder^10^.

Next generation sequencing (NGS) technologies have made it possible to study whole genomes or exomes at a single base pair resolution. In disease studies, trio-based sequencing involves whole exome sequencing (WES) or whole genome sequencing (WGS) of an affected offspring and their parents. This approach allows for the identification of genetic variants unique to the affected offspring and is commonly used to study sporadic genetic disorders caused by DNMs^11–17^.

In standard approaches, DNMs are detected by first inferring the genotypes of each individual in a trio separately using standard variant calling algorithms such as Genome Analysis Toolkit (GATK)^18^ or Samtools^19^. The putative DNMs are then identified by comparing the genotypes of the offspring to those of the parents. However, this approach is inefficient and yields many false positives. The error rates in NGS data are much higher than the expected DNM rate, and any miscalled genotypes in either the parents or the offspring will result in false positives. In recent years, many computational methods have been developed specifically for calling DNMs from NGS data. These methods use the raw read-alignment data or genotype likelihoods for each possible genotype to determine a DNM call. Four widely-used DNM calling tools are DeNovoGear^20^, VarScan 2^21^, TrioDeNovo^22^ and PhaseByTransmission^23^.

Independent benchmarking and evaluation of genomics software, such as variant callers, is critical for establishing best practices. Several studies have compared variant calling pipelines for detecting SNVs and insertions/deletions (indels)^24–26^, including somatic mutations^27–29^, across different sequencing technologies, but were not focused on DNMs which can only be determined by using family trio data. One prior study evaluated DNM callers in humans^30^ using WES data from the Ashkenazi Jewish trio set (NA12878). However, this comparison did not include widely used DNM callers such as TrioDeNovo, DeNovoGear and PhaseByTransmission. Additionally, the evaluation of this study did not incorporate any validated DNMs. Instead, GC (guanine-cytosine) content, substitution type and single nucleotide variant (SNV) density were used as proxy evaluation metrics.

In this study, we present a systematic evaluation of commonly used DNM calling tools, as well as a more recent deep convolutional neural network (CNN) DNM caller (DeNovoCNN)^31^ for WGS trio data (Fig. 1). Our assessment incorporates a dataset comprising validated DNMs as well as a simulated trio dataset generated in-house. *De novo* single nucleotide variants (*dn*SNVs) and *de novo* insertions/deletions (*dn*INDELs) are compared separately for characteristics affecting DNM detection, such as read depth and allelic fraction. Our objective is to offer recommendations of best practices for the identification of DNMs from WGS trio data.

**Figure 1.**
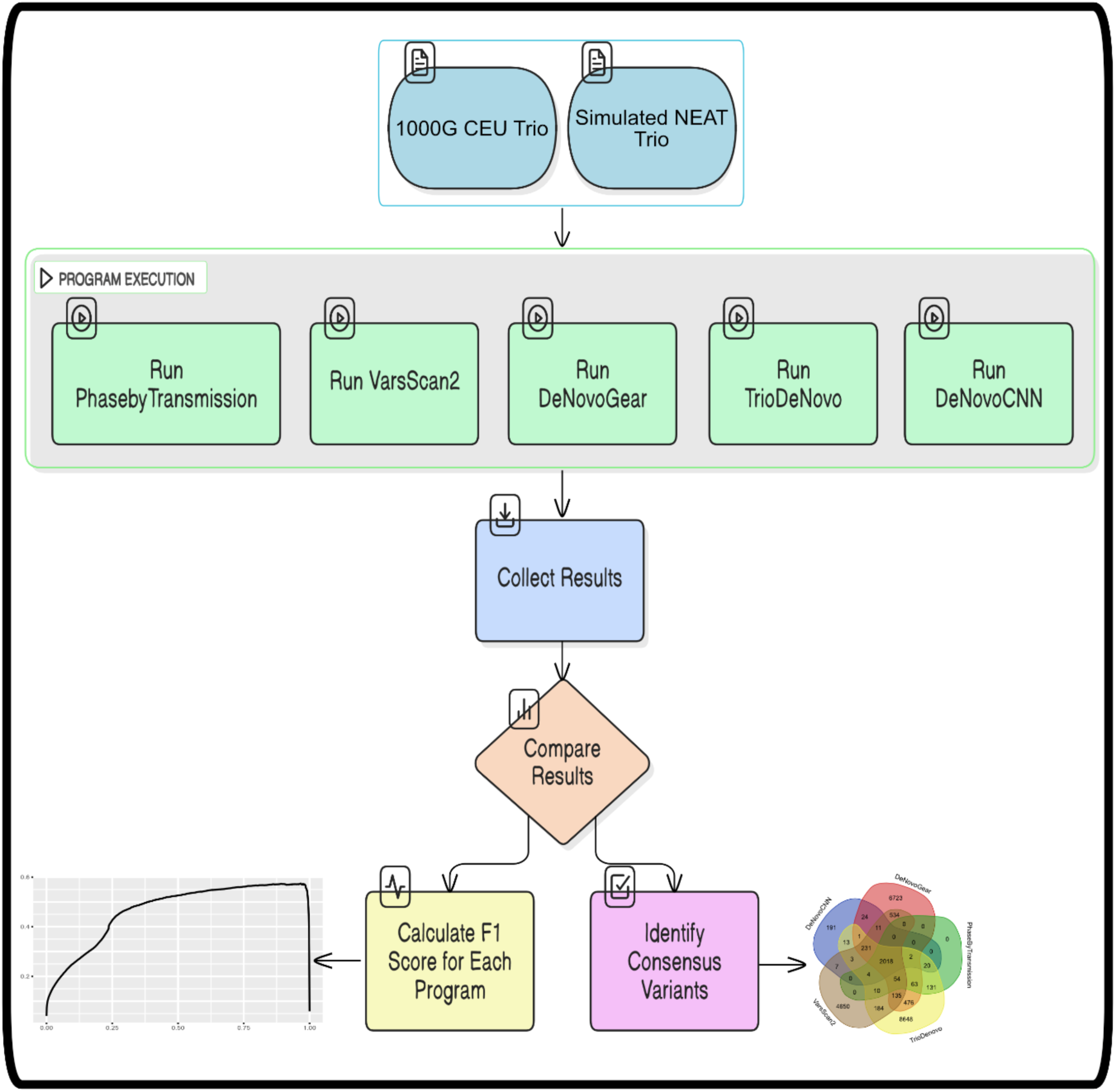
Workflow for benchmarking de novo mutation (DNM) callers. Overview of the analysis pipeline applied to the 1000 Genomes CEU trio and the simulated WGS NEAT trio datasets. Both datasets were processed through five DNM callers: PhaseByTransmission, VarScan2, DeNovoGear, TrioDeNovo and DeNovoCNN. Performance was evaluated using F1 scores based on true positive DNMs identified in each dataset. Additionally, consensus calls across the five tools were examined.

## MATERIAL AND METHODS

### Real dataset

We used the Illumina HiSeq WGS data of one CEU (Central European from Utah) trio from the 1000 Genomes Project (1000G) for evaluation^32^. The trio samples we used were NA12891 (father), NA12892 (mother), and NA12878 (female offspring). Their aligned CRAM files (based on the GRCh38 reference genome) were downloaded from ftp://ftp.1000genomes.ebi.ac.uk/vol1/ftp/data_collections/illumina_platinum_pedigree/data/CEU/NA12878/ and were converted to BAM files for further bioinformatics processing. The depth of coverage of BAM files for father, mother, and offspring was 60x, 60x, and 66x, respectively. The genome co- ordinates of variants were lifted over from hg19 to hg38 using UCSC genome browser’s hgLiftOver feature^33^. The dataset contains experimentally validated DNMs, alongside other variant types.

Specifically, the dataset comprises 48 germline DNMs, 888 cell line somatic DNMs, 125 inherited polymorphisms and 1,256 false positive DNMs, all originating from autosomal chromosomes. For the DNM variant caller comparison, the combined germline and cell line somatic DNMs (936) will be used as positive DNMs. A Variant Call Format (VCF) containing all samples in the trio was generated by executing the best practices standard variant calling pipeline of GATK (version 4.0)^37^ using default parameters.

### Simulated dataset

Using an in-house application called TrioSim, we generated a synthetic WGS-based family trio set of BAMs (Fig. S1). TrioSim utilises SIMdrom^34^ to generate mother and father VCFs with allele frequencies observed in the European population data available in the 1000G project. The two resulting VCF files were merged to create one parental VCF file encompassing variants from both parents. The offspring VCF file was then generated by applying the Mendelian inheritance law to the merged parental VCF file, to infer the genotypes of the offspring. Next, 100 DNMs were randomly sampled from the public database denovo-db^35^ and spiked into the offspring VCF file. These DNMs comprised 92 *dn*SNVs and 8 *dn*INDELs. All simulated and spiked-in variants originated from autosomal chromosomes, consistent with real data. Lastly, Illumina-based NGS reads were simulated using the NEAT^36^ open-source software package with the GRCh38 reference genome, at 30x coverage depth. The final output of TrioSim includes three independent BAM files representing the trio (mother, father, and offspring). TrioSim is available at https://github.com/VCCRI/TrioSim

To ensure data integrity and quality, the generated BAM files underwent post-processing procedures in adherence to GATK’s best practices guidelines. A VCF file utilising the simulated trio’s generated BAMs was created using GATK (version 4.0), with default parameters. The VCF file was checked for Mendelian inconsistencies using the +mendelian plug-in provided by bcftools^19^.

### *De novo* mutation calling

#### DeNovoGear

DeNovoGear^20^ is a software application that detects DNMs from familial and somatic tissue using NGS trio data. Its model consists of individual genotype likelihoods, transmission probabilities and prior probabilities to model the likelihood of observing a polymorphism or a DNM at any given site (Table 1). Users can specify the prior probability of observing a DNM which can be used to increase or decrease calling sensitivity. It employs beta-binomial distributions that are fitted to alternate read frequencies for both SNPs and indels to facilitate mutation detection. The “dng dnm” subcommand of DeNovoGear was executed to detect DNMs in trios. DeNovoGear required as inputs a Pedigree (PED) format file and a trio based Binary Variant Call Format (BCF) file. The output is a single text file containing a row for each putative DNM, with various fields including event type (Point Mutation/INDEL), physical location and the posterior probability of the most likely *de novo* genotype configuration.

**Table 1.** Features of DNM calling tools.

|  | **DeNovoGear** | **VarScan 2** | **TrioDeNovo** | **PhaseByTransmission** | **DeNovoCNN** |
| --- | --- | --- | --- | --- | --- |
| **Method**<br>**/Algorithm** | Beta-binomial based likelihood model and evaluation of joint trio likelihood. | Statistical significance test for variant calling and heuristics. | Bayesian model selection framework using individual genotype likelihoods. | Bayesian framework using individual genotype likelihoods. | Deep convolutional neural network that encodes the alignment of sequence reads for a trio images |
| **Input data** | BCF (mpileup) and PED files | BCF (mpileup) | VCF file | VCF file | BAM/CRAM and VCF files |
| **Output data** | Text file | VCF files | VCF file | VCF file | Text file |
| **Output**<br>**Confidence**<br>**score for DNM** | Posterior probability of the most likely *de novo* genotype configuration (PPDM) | Phred-scaled P-value from Fisher's exact test | Bayes Factor (DeNovo Quality) | Phred-scaled posterior probability (Phred Score) | DeNovoCNN Probability |
| **Phasing**<br>**information** | Yes | No | No | Yes | No |

#### VarScan 2

VarScan2^21^ is a variant calling tool that detects somatic and germline mutations from NGS trio data^21^. The DNM calling “trio” subcommand of VarScan 2 detects high confidence DNMs by leveraging the familial relationship present in trio datasets, thereby identifying apparent Mendelian Inheritance Errors (MIEs) (Table 1). It first calls all variants across all three samples using the standard way of mpileup2snp, using default values for parameters “--min-var-freq” and “--p-value”. Subsequently, it identifies any variants with apparent MIEs, i.e., variants that are present in the offspring but absent in parental samples. In such circumstances, it re-calls the samples with relaxed parameters and reclassifies with some evidence (--adj-var-freq and --adj-p-value). Often, this corrects MIEs, and corrected genotypes are reported. If not, the site is reported as “mendelError” (in the “FILTER” field) and/or “DENOVO” (in the “INFO” field). The VarScan 2 “trio” subcommand utilises a trio BCF file as input and produces two output VCF files: one for SNVs and another for indels. The output VCF files contain “INFO” fields for all called SNVs and indels. If a DNM is called, the “STATUS VCF” field will contain a value of “3”. For high confidence DNM calls, the variant is also annotated with the text label “DENOVO”.

#### TrioDeNovo

TrioDeNovo^22^ is a program which uses a Bayesian framework to detect DNMs from NGS trio data without specifying a prior mutation rate^2^ (Table 1). Firstly, it evaluates the evidence of DNMs using trio data and then adjusts the effect of mutation rates on DNM calling through prior odds. For input, the user provides a PED file, and a VCF file with PL (Phred-scaled genotype likelihood) or GL (genotype likelihood) fields. The output is a VCF file containing the detected DNMs.

#### PhaseByTransmission

PhaseByTransmission^23^ is a software module integrated with GATK (version 3.8) to detect DNMs from NGS trio data. By utilising genotype likelihoods and a mutation prior, it calculates the most likely genotype combination and phases trios and parent/offspring pairs (Table 1). The software requires a PED file and a VCF file as input, and it outputs a VCF file and a Mendelian violations file. PhaseByTransmission computes genotype combinations of the trio (i.e. mother-father-offspring) and supports bi-allelic cases, with 27 possible genotype-trio combinations. For our comparison, we have used the trio genotype combination of “AA-AA-AB” (i.e. mother-father-offspring), where the A is the reference allele. PhaseByTransmission is included in the GATK software, version 3.8 release.

#### DeNovoCNN

DenovoCNN^31^ is a variant calling tool that utilises an image classification deep convolutional neural network (CNN) to identify DNMs in a family trio. It converts NGS sequence reads for a trio into 160×164 resolution images, which are then processed by the CNN as a vision classification task to identify the images as DNMs or inherited variants (Table 1). DenovoCNN takes as input the trio’s individual VCF and BAM/CRAM files to output a prediction csv file and a text file containing the list of variants. The prediction csv file contains the DeNovoCNN probability for each variant evaluated, with a range of 0 (lowest probability for DNM) – 1 (highest probability for DNM), as well as read coverage for each member of the trio. DeNovoCNN suggests a probability cutoff of 0.5 and above for good DNM candidates.

### Performance Evaluation

To evaluate the performance of the five DNM callers in detecting DNMs, we generated confusion matrices and calculated F1 scores. For each caller, we obtained results using two methods: DNMs called using the program’s default setting, and DNMs based on a “benchmarking-set maximal F1 threshold”. We determined the “benchmarking-set maximal F1 threshold” for each caller by calculating F1 scores across the range of possible thresholds (the range of each caller’s confidence scores) and identifying the threshold that resulted in the highest F1 score (Fig. S2). Results using the “benchmarking-set maximal F1 threshold” represent the best performance for each caller, for the dataset. This enabled a fair comparison of results between the five callers.

DeNovoGear and PhaseByTransmission provide phased genotype information, which can be used to infer haplotypes from genotype data. However, as TrioDeNovo, DeNovoCNN and PhaseByTransmission do not provide this information, in our analysis we only considered unphased genotype information in comparing the DNM callers.

## RESULTS

We evaluated the DNM calling methods DeNovoGear, VarScan 2, TrioDeNovo, PhaseByTransmission and DeNovoCNN using both real (1000G CEU) and simulated datasets of WGS trios.

### Real dataset

PhaseByTransmission detected the largest number of DNMs (60,079), a large portion being *dn*INDELs (44,463), while DeNovoCNN detected the lowest number of DNMs (4,069) from the 1000G CEU trio dataset (Table 2). In evaluating the results, we found that some DNMs detected by the callers had missing genotypes for one or both parents. To remove these falsely classified DNMs, we applied genotype-based filters using the genotype pattern of “0/0” (homozygous for the reference allele) for the mother and father, and “0/1” (heterozygous for the alternative allele) for the offspring. After filtering, PhaseByTransmission had the lowest number of DNM calls for *dn*SNVs (2,302), followed by DeNovoCNN (2,525), VarScan 2 (7,841) and DeNovoGear (10,272) (Table 3).DeNovoCNN (1,544),and TrioDeNovo (9,655) detected the lowest and highest number of *dn*INDELs, respectively.

**Table 2.**
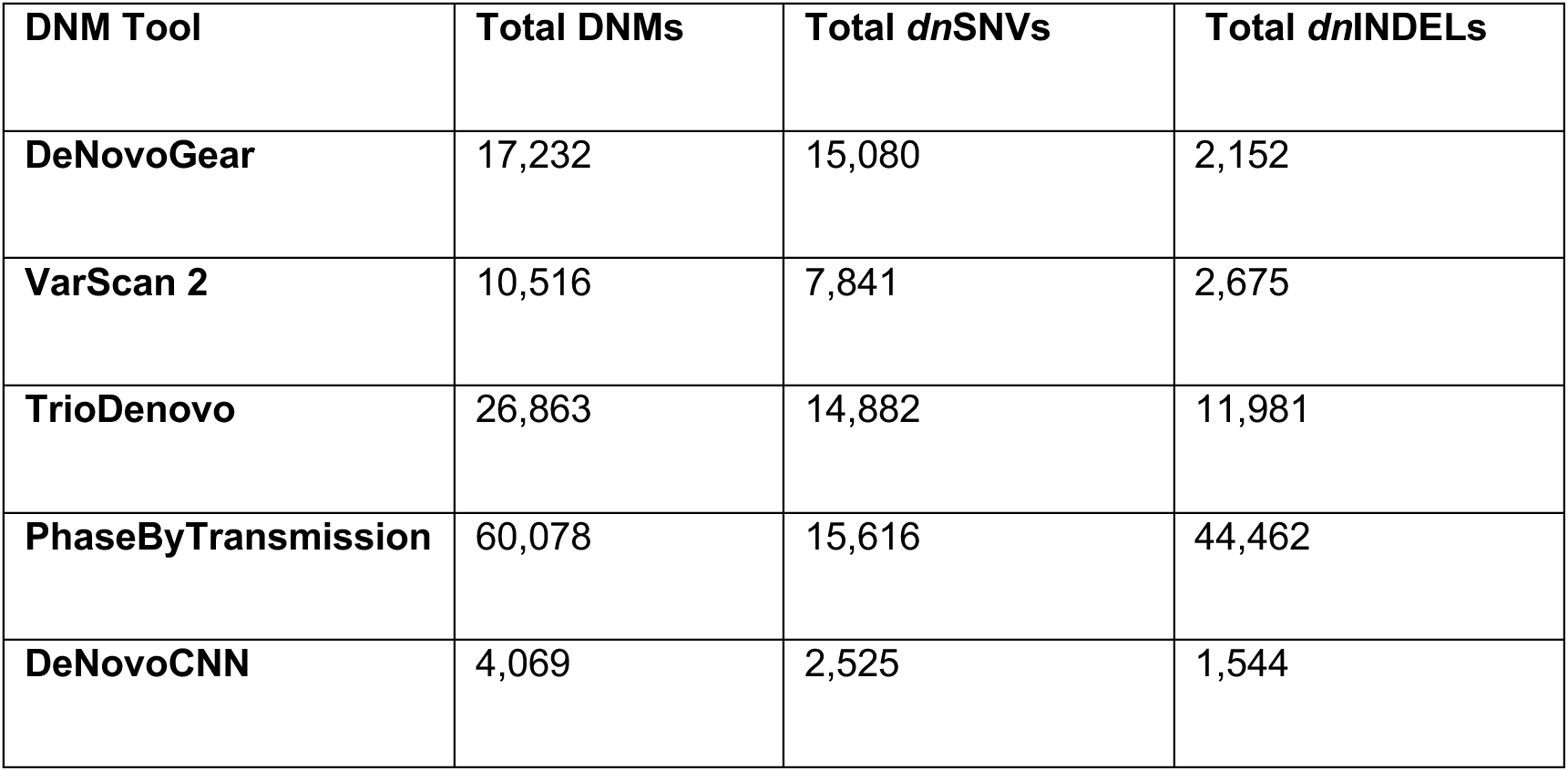
Total counts of DNMs, *dn*SNVs and *dn*INDELs called by each DNM caller using the 1000G CEU trio, using default caller settings.

**Table 3.** Total counts of DNMs, *dn*SNVs and *dn*INDELs called by each DNM caller using the 1000G CEU trio, following genotype filtering. Genotype filters were applied against the callers’ results from the default setting run. (Father, Mother, Offspring: 0/0, 0/0, 0/1).

| **DNM Tool** | **Total DNMs** | **Total *dn*SNVs** | **Total *dn*INDELs** |
| --- | --- | --- | --- |
| **DeNovoGear** | 11,558 | 10,272 | 1,286 |
| **VarScan 2** | 10,516 | 7,841 | 2,675 |
| **TrioDeNovo** | 21,648 | 11,993 | 9,655 |
| **PhaseByTransmission** | 4,926 | 2,302 | 2,624 |
| **DeNovoCNN** | 4,069 | 2,525 | 1,544 |

We investigated the concordance of DNMs called by the five DNM callers for *dn*SNVs (Fig. 2a) and *dn*INDELs (Fig. 2b). The concordance was extremely low, with only 2,018 *dn*SNVs (8.4%) and 111 *dn*INDELs (0.8%) called by all callers. TrioDeNovo gave the largest number of unique calls (not identified by any other callers), with 8,648 *dn*SNVs and 6,817 *dn*INDELs. In contrast, PhaseByTransmission had no unique *dn*SNVs calls and only 7 unique *dn*INDELs.

**Figure 2.**
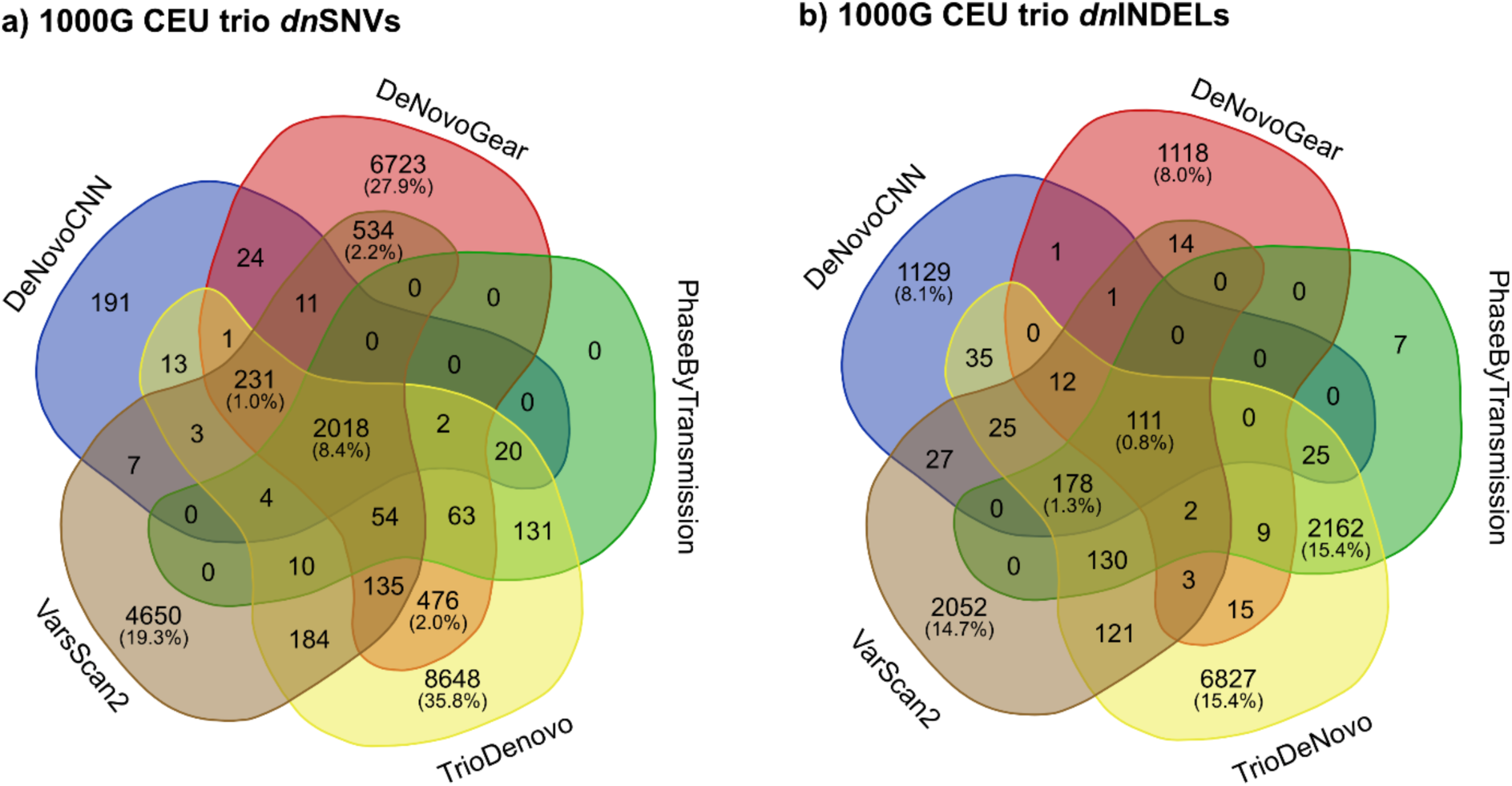
Concordance between DNM callers on the 1000G CEU trio dataset. The Venn diagrams show counts of DNMs called by combinations of the 5 DNM callers for (a) *dn*SNVs and (b) *dn*INDELs. The numbers in parentheses indicate the percentages of total DNMs. Percentages below 1% are only shown for DNMs identified by all five methods.

Using the DNM callers’ default score thresholds, TrioDeNovo called the highest number of true positive calls (929 TPs) (Table 4). However, it also produced the highest number of false positive calls (FPs=11,064) and had the lowest F1 score (0.14371). In comparison, PhaseByTransmission called a slightly lower number of TPs (918), but it called the lowest number of FPs (1,384) and achieved the highest F1 score (0.56702).

**Table 4.** Performance results using default DNM caller thresholds for 1000G CEU trio dataset. The five callers were evaluated on the 1000G CEU trio *dn*SNVs. The following metrics were calculated: True Positives (TP), False Negatives (FN), False Positives (FP), and F1 score.

| DNM caller | TP | FN | FP | F1 score |
| --- | --- | --- | --- | --- |
| DeNovoGear | 921 | 15 | 9,351 | 0.16435 |
| VarScan 2 | 916 | 20 | 6,925 | 0.20873 |
| TrioDeNovo | 929 | 7 | 11,064 | 0.14371 |
| PhaseByTransmission | 918 | 18 | 1,384 | 0.56702 |
| DeNovoCNN | 910 | 26 | 1,615 | 0.52586 |

We calculated the “benchmarking-set maximal F1 threshold” for each DNM caller (see Methods – Performance evaluation). Using “benchmarking-set maximal F1 thresholds”, DeNovoGear achieved the highest F1 score (0.58887), and substantially reduced its FPs (1,168) (Table 5).

**Table 5.**
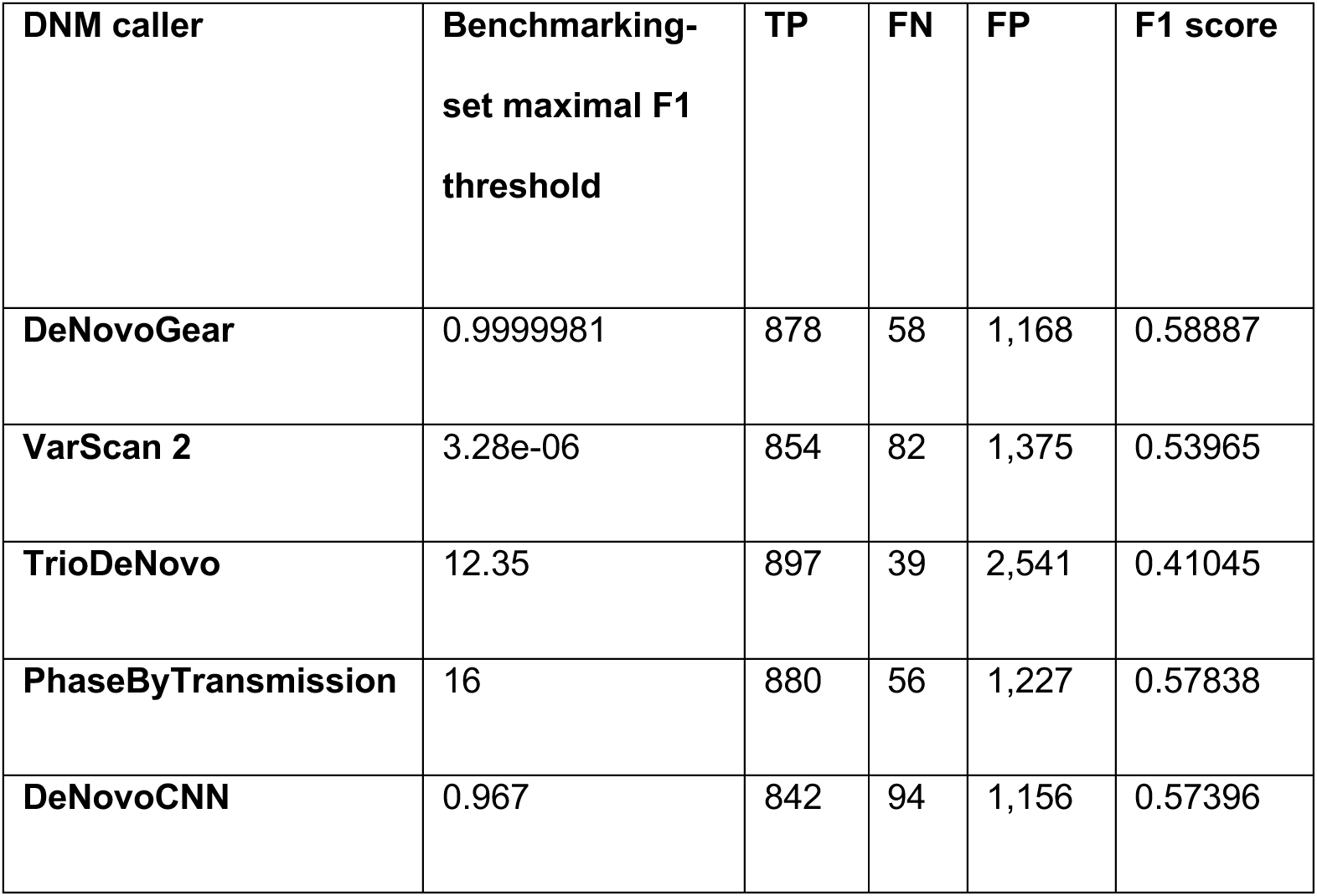
Performance results using “benchmarking-set maximal F1 thresholds” for 1000G CEU trio dataset. The five callers were evaluated on the f 1000G CEU trio *dn*SNVs. The following metrics were calculated: True Positives (TP), False Negatives (FN), False Positives (FP) and F1 score.

| DNM caller | Benchmarking-set maximal F1 threshold | TP | FN | FP | F1 score |
| --- | --- | --- | --- | --- | --- |
| DeNovoGear | 0.9999981 | 878 | 58 | 1,168 | 0.58887 |
| VarScan 2 | 3.28e-06 | 854 | 82 | 1,375 | 0.53965 |
| TrioDeNovo | 12.35 | 897 | 39 | 2,541 | 0.41045 |
| PhaseByTransmission | 16 | 880 | 56 | 1,227 | 0.57838 |
| DeNovoCNN | 0.967 | 842 | 94 | 1,156 | 0.57396 |

PhaseByTransmission and DeNovoCNN were very close behind, with F1 scores of 0.57838 and 0.57396, respectively, but DeNovoCNN had the lowest TP value at 842 out of 936. VarScan 2 followed with an F1 score of 0.53965, and TrioDeNovo achieved an F1 score of 0.41015, producing the highest number of FPs (2,541).

### Properties of True Positive and False Positive DNMs identified by DNM callers

The accuracy of each algorithm in detecting DNMs depends on both the sequencing coverage and the true variant allele frequency (VAF) of the mutations. We investigated the relationship between read depth at the DNM positions and the estimated VAF of DNMs identified by all callers. To maintain consistent values between the DNM callers, VAF was calculated using allelic counts obtained using the GATK tool “CollectAllelicCounts”. For true positive *dn*SNVs, all the DNM callers had similar VAFs with a combined mean value of 0.45 and standard deviation of 0.08 and were identified in positions with read depths between 27 and 80 (Fig. 3a).

**Figure 3.**
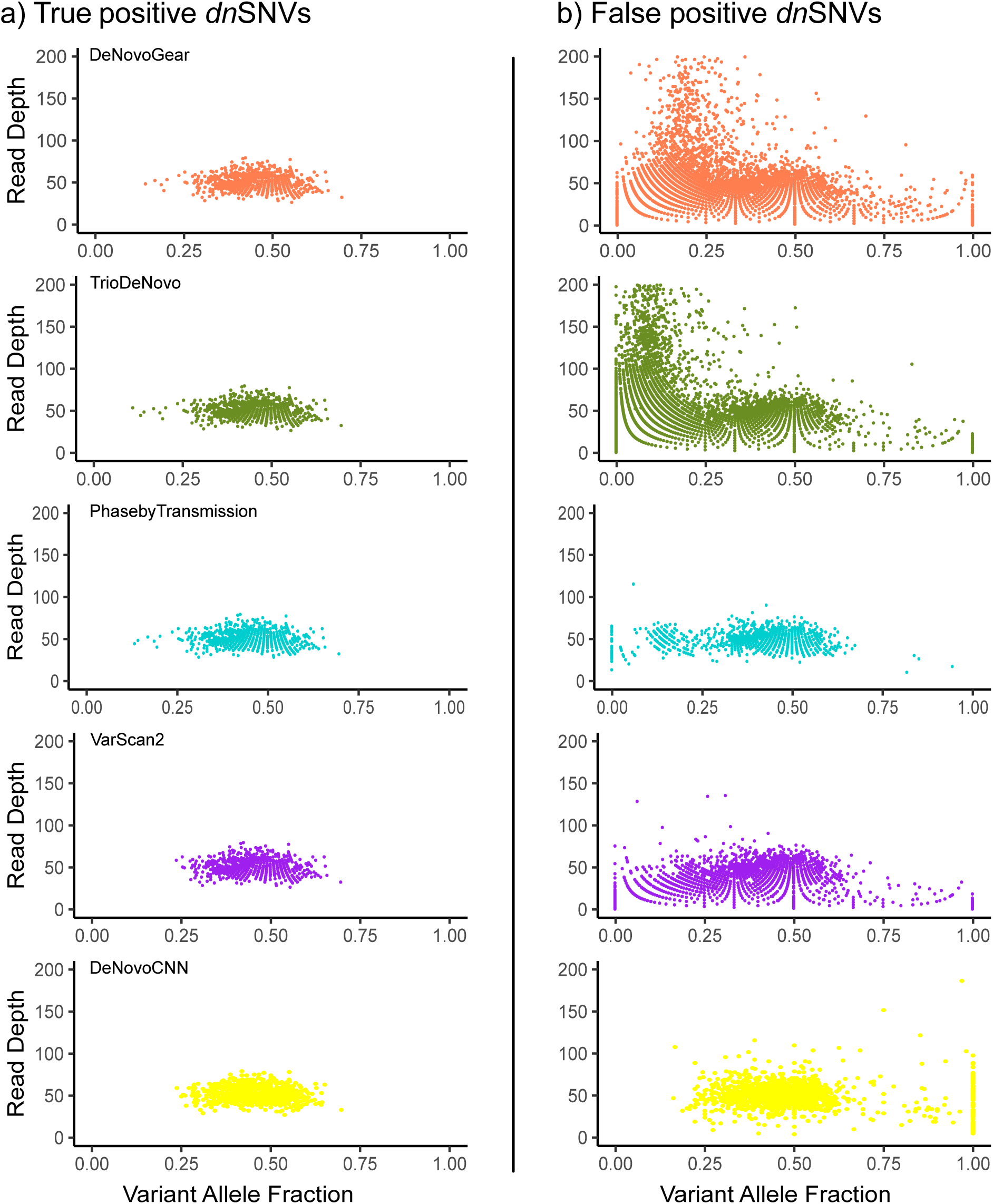
Variant Allele Fraction vs. Read Depth for *dn*SNVs in proband (NA12878) for each DNM caller using CEU trio from 1000G. a) True positives and b) false positives for each DNM caller are shown separately. The y-axis is cropped at 200.

DeNovoGear and TrioDeNovo identified false positive *dn*SNVs across a wide range of VAFs and read depths, especially at lower VAF levels when the read depth was high (>100 reads) (Fig. 3b). By comparison, DeNovoCNN, PhaseByTransmission and VarScan 2 did not detect false positive *dn*SNVs at high read depths. DeNovoCNN did not produce any false positive dnSNVs in the low VAF range (< 0.2), whereas the other callers generated false positive *dn*SNVs across the full range of VAFs (0-1). We note that the DNM callers can use different methods of filtering lower quality reads for use in the VAF calculation (compared to “CollectAllelicCounts”), which may explain some discrepancies, for example the presence of DNM calls with VAF values of 0. There were no true positive *dn*INDELs called, so only false positive *dn*INDELs were analysed. For false positive *dn*INDELs, we observed that all callers identified *dn*INDELs with lower VAF. Both TrioDeNovo and DeNovoCNN had false positive *dn*INDEL calls across the full VAF range of 0-1 (Fig. S3).

We also investigated the VAF distribution within the trio samples for both true positive and false positive DNMs, as a factor influencing whether DNMs are called. For true positive *dn*SNVs, we observed that the parent’s VAFs in the DNM positions were extremely low or zero (father’s mean VAF = 0.0002 and standard deviation = 0.002, mother’s mean VAF = 0.0001 and standard deviation = 0.001, across all callers) (Fig. 4a). In contrast, the offspring’s VAF were centred around 0.5, consistent with the expected profile of a heterozygous DNM.

**Figure 4.**
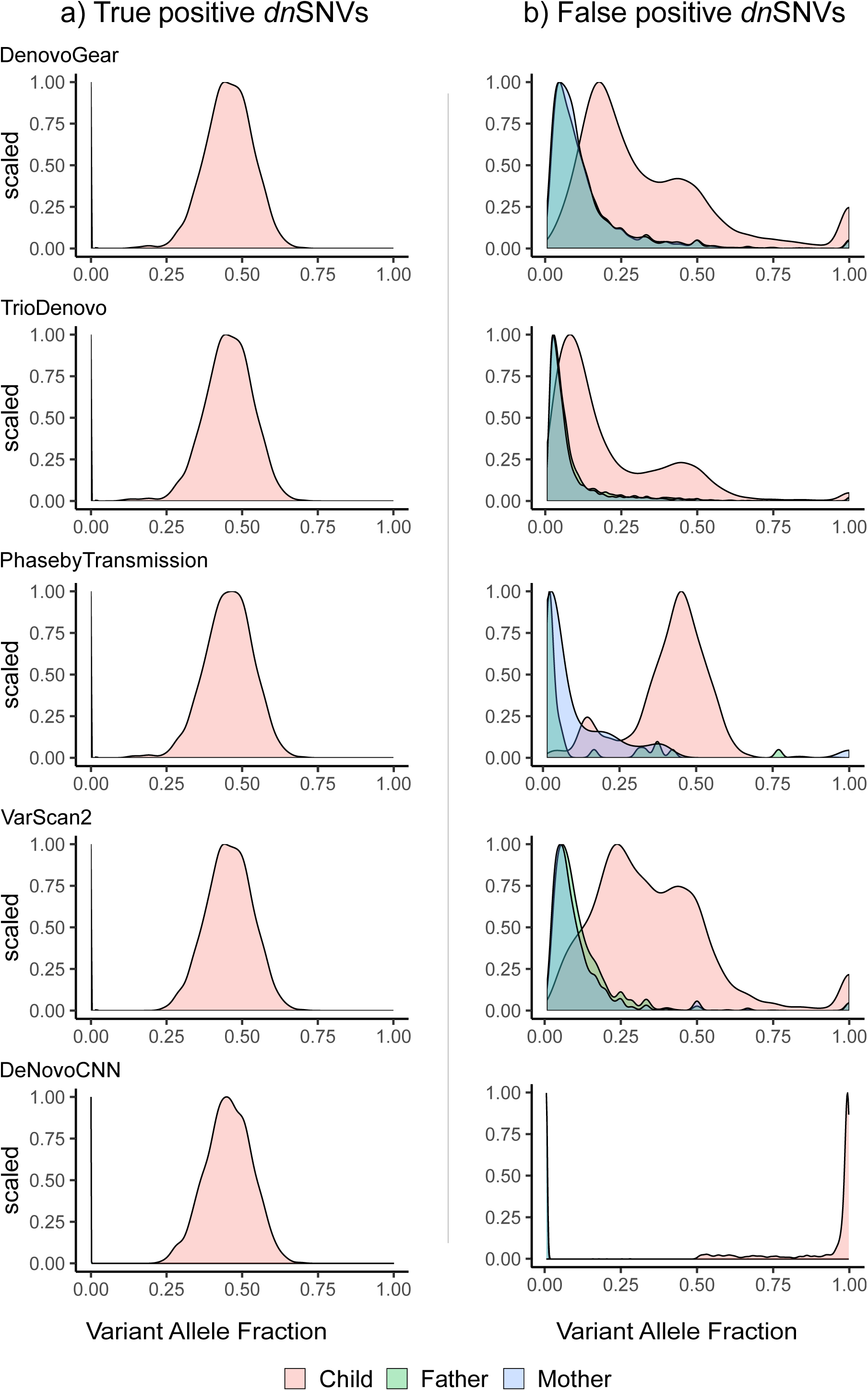
Densities of Variant Allele Fraction of *dn*SNVs in the 1000G CEU trio. VAF density for. a) True positive *dn*SNVs and b) False positive *dn*SNVs for each DNM caller are shown separately. In the true positives plots for all DNM callers and the DeNovoCNN false positives plots, the VAF distributions for the parents are visible only as a spike close to VAF=0.

In the case of false positive *dn*SNVs, we observed that for all callers, the parental VAFs were lower, but not as extremely low or zero as those seen in true positive dnSNVs - except for DeNovoCNN, which showed parental VAFs that closely mirrored the extreme low or zero levels observed in true positives (Fig. 4b). The offspring VAFs were spread across the VAF range for DeNovoGear, TrioDeNovo and VarScan2, but PhaseByTransmission, maintained a more likely *dn*SNV offspring profile of a VAF value centred at 0.5. For DeNovoCNN, the majority of the offspring’s false positive *dn*SNVs VAF were close to 1. The variation in VAF in the offspring indicates that some of these false positive *dn*SNVs may either be systematic errors present across the trio due to low read quality, orinherited variants observed with lower VAFs in the parents. For false positive *dn*INDELs, we observed a similarly large concentration of very low VAFs for the mother and father (Fig. S4). For the offspring, all DNM callers except DeNovoCNN showed low VAFs, whereas DeNovoCNN had offspring VAF across the full range from 0 to 1.

We further investigated the relationship between VAF and read depth using the consensus of different numbers of callers for *dn*SNVs (Fig. S5). Out of 936 validated true positive *dn*SNVs, 895 were detected by all five DNM callers. These 895 dnSNVs had VAFs ranging between 0.24 and 0.70 and a mean read depth of 51. The false positive *dn*SNVs called by all five DNM callers also had very similar VAF distributions, ranging between 0.2 to 0.94, and a mean read depth of 50. As the number of consensus DNM callers decreased, the VAF range increased.

We also investigated patterns in the output confidence scores of TP and FP DNMs for each caller (Fig. 5). The scores for DeNovoGear and VarScan 2 showed a clear separation between TP and FP DNMs, indicating better calibration. DeNovoGear uses posterior probability for its DNM score (PPDM).

**Figure 5.**
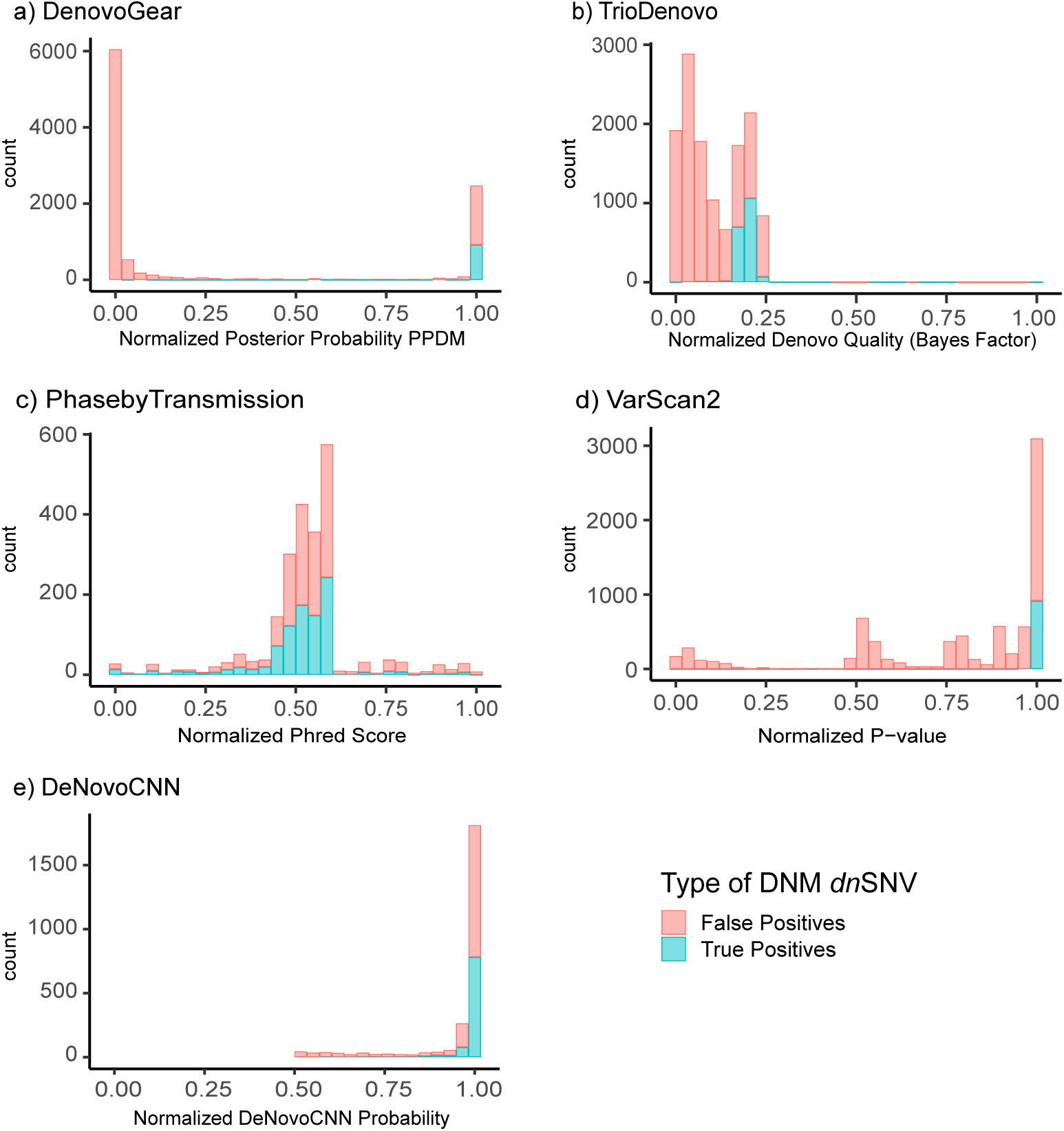
Distribution of the DNM scores for each DNM caller using the 1000G CEU trio. True positive and False positive *dn*SNVs are shown for each DNM caller. The DNM caller’s raw confidence score for each variant has been normalized for plotting purposes. (a) Posterior Probability of *de novo* mutation (PPDM) by DeNovoGear; (b) Denovo Quality log 10 (Bayes Factor) by TrioDeNovo ;(c) P-value by VarScan 2; (d) Phred score by PhaseByTransmission; and e) DeNovoCNN Probability by DeNovoCNN.

The majority (916/921) of TP DNMs called by DeNovoGear had PPDM > 0.99, while 83.5% (7809/9351) of FP DNMs had PPDM < 0.99. For VarScan 2, which uses a normalised P-value, all 916 TP DNMs had p-value < 9.4048 x 10^-4^, while 84.5% of FP DNMs had p-value > 0.05. TrioDeNovo, which uses a normalised Denovo Quality score, showed overlap between TP and FP DNMs, though TPs were concentrated in the higher quality score range. For PhaseByTransmission, which uses a normalised Phred score, there was no clear distinction between the scores of TP and FP DNMs. DeNovoCNN also show no clear distinction between TP and FP DNMs, as both were identified across most of its confidence score range – from its maximum value of 1 down to just above its recommended cutoff of 0.5.

### Simulated Datasets

We simulated trio WGS data with 100 spiked-in DNMs using a custom framework, TrioSim, as described in the methods section (Figure S1). Of these 100 spiked-in DNMs, 92 were *dn*SNVs and 8 were *dn*INDELs. We applied the five DNM callers to the simulated data and evaluated their performance.

Using the default settings for each caller, DeNovoGear called the largest number of DNMs (1,431), followed by TrioDeNovo, VarScan 2, PhaseByTransmission and finally DeNovoCNN, which called only 281 DNMs (Table 6). DeNovoGear also identified the most *dn*SNVs (1,391), while PhaseByTransmission identified the fewest (117). TrioDeNovo detected the largest number of *dn*INDELs (824), while DeNovoGear detected the smallest number (40). DeNovoGear, VarScan 2, DeNovoCNN and TrioDeNovo successfully identified all 92 TP *dn*SNVs, while PhaseByTransmission identified only 35 of them. Due to the high numbers of *dn*SNVs called, DeNovoGear reported the greatest number of false positives (1,299), followed by VarScan 2 (584), TrioDeNovo (434), DeNovoCNN (142) and PhaseByTransmission (82).

**Table 6.** Total DNMs, *dn*SNVs, *dn*INDELs, True Positives (TP) and False Positives (FP) for each DNM caller using NEAT simulated data using default DNM caller settings.

| **DNM Tool** | **Total DNMs** | **Total *dn*SNVs** | **Total *dn*INDELs** | **TP *dn*SNVs** | **FP *dn*SNVs** |
| --- | --- | --- | --- | --- | --- |
| **DeNovoGear** | 1,431 | 1,391 | 40 | 92 | 1,299 |
| **VarScan 2** | 759 | 676 | 83 | 92 | 584 |
| **TrioDeNovo** | 1,350 | 526 | 824 | 92 | 434 |
| **PhaseByTransmission** | 315 | 117 | 198 | 35 | 82 |
| **DeNovoCNN** | 281 | 234 | 47 | 92 | 142 |

We calculated concordance and evaluation metrics for *dn*SNVs following the same procedure as with the real 1000G CEU trio dataset. We found that the concordance between the 5 callers was exceptionally low at 3.9% (Fig. 6). This low concordance was consistent with the results observed from the real dataset. DeNovoGear had the highest number of unique DNM calls (815).

**Figure 6.**
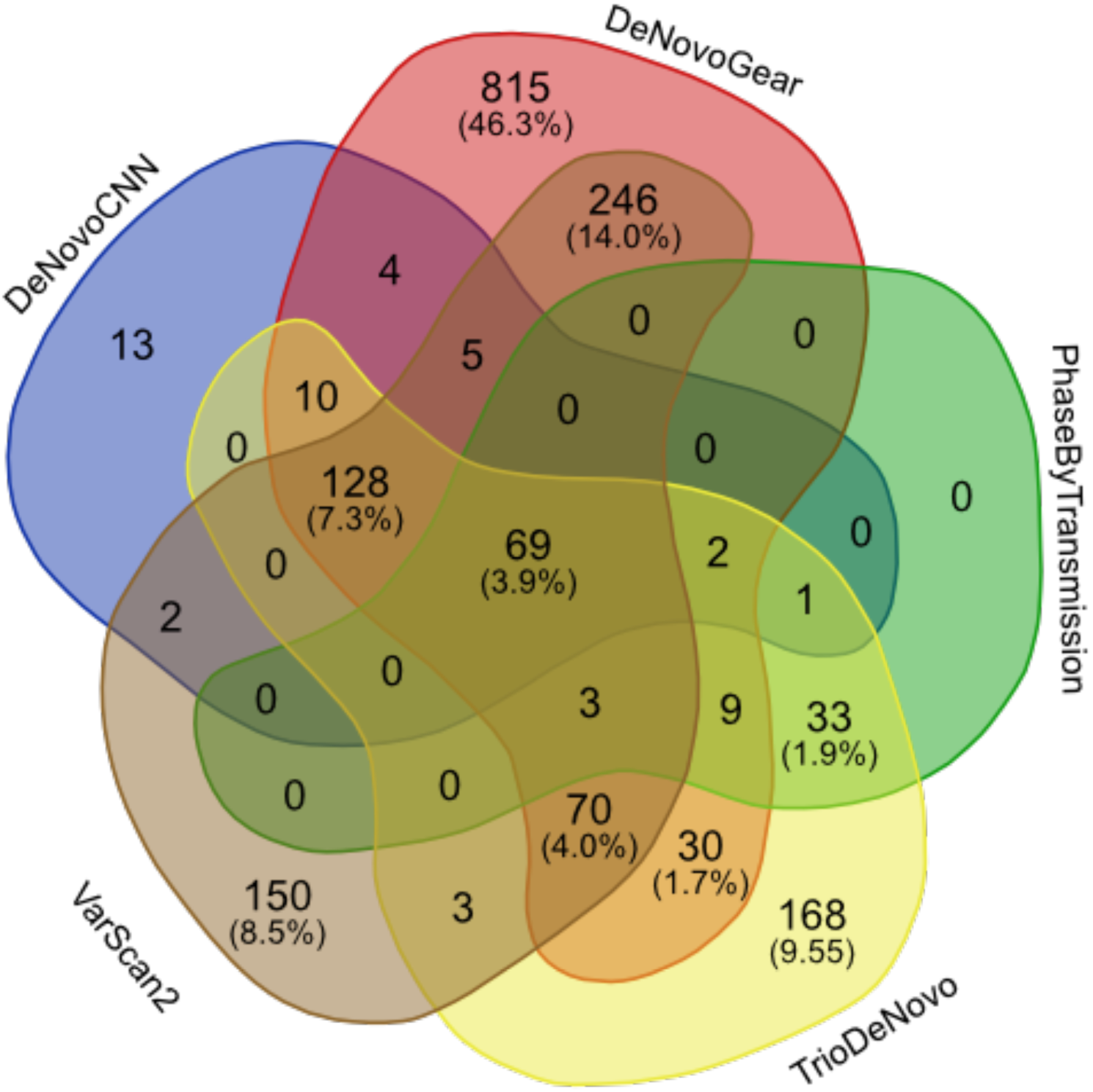
Concordance between DNM callers on the NEAT-simulated data. The Venn diagram illustrates total counts for *dn*SNVs called by each DNM caller. The numbers in the parentheses indicate the percentages of total *dn*SNVs. Percentages below 1% are not shown.

For performance evaluation, we utilised the same “benchmarking-set maximal F1 threshold” procedure as for the real dataset for the callers (see Methods – Performance evaluation). DeNovoCNN achieved the highest F1 score of 0.610687, followed by DeNovoGear with 0.56089 (Table 7). Next was TrioDeNovo with an F1 score of 0.42703, followed by PhasebyTransmission at 0.33493. Finally, VarScan 2 had the lowest F1 score of 0.23349, mainly due to its high number of FPs (483).

**Table 7.** Performance results using “benchmarking-set maximal F1 thresholds” on the NEAT simulated dataset. The following metrics were calculated: True Positives (TP), False Negatives (FN), False Positives (FP), and F1 score.

| **DNM caller** | **Benchmarking-set maximal F1 threshold** | **TP** | **FN** | **FP** | **F1 score** |
| --- | --- | --- | --- | --- | --- |
| DeNovoGear | 0.99102 | 76 | 16 | 103 | 0.56089 |
| VarScan 2 | 0.00012 | 76 | 16 | 483 | 0.23349 |
| TrioDeNovo | 9.32 | 79 | 13 | 199 | 0.42703 |
| PhaseByTransmission | 1 | 35 | 57 | 82 | 0.33493 |
| DeNovoCNN | 0.961 | 80 | 12 | 90 | 0.610687 |

## DISCUSSION

In this study, we present a systematic evaluation of four widely used DNM calling tools: DeNovoGear, VarScan 2, TrioDeNovo, and PhaseByTransmission, and a more recent tool, DeNovoCNN. We developed a software framework for generating simulated NGS trio datasets named TrioSim, which is freely available. Using a simulated dataset, as well as an experimentally validated 1000G CEU trio dataset, we investigated the performance of each caller.

We found that the raw number of calls varied dramatically between the different callers and the concordance among the five callers was strikingly low, with 8.4% for *dn*SNVs and 0.8% for *dn*INDELs in the 1000G CEU dataset. For both the real and simulated datasets, all DNM callers showed high sensitivity as they were able to detect the majority of true DNMs.

When detecting DNMs from NGS data, the large scale of WGS data eclipses the small number of true DNMs present. In this setting, a tendency to call FPs is particularly costly as it can lead to overwhelmingly large numbers of (mostly false positive) DNM calls. To provide a more realistic classification setting, we designed evaluation datasets with significant imbalances in favour of negative variants. In this imbalanced setting, F1 score was chosen as an appropriate evaluation metric.

We provide performance results using each caller’s default thresholds, as we expect that many users would adopt the default settings, so performance at these thresholds is of interest. We also provide results using “benchmarking-set maximal F1 thresholds”, which allows a fairer comparison of each caller’s ability to identify DNMs. Overall, we identified DeNovoGear and DeNovoCNN as the best performing DNM callers. DeNovoGear achieved the highest F1 score on the real 1000G dataset, while DeNovoCNN had the highest F1 score on the simulated data.

We found that F1 scores were relatively low for all callers, with the highest F1 scores achieved on the real and simulated datasets being 0.58887 and 0.610687, respectively. The large numbers of false positive DNMs is a major issue for all the callers, with some callers having as much as 10x as many false positive DNMs as compared to true positive DNMs. Taken together, these results are indicative of significant performance issues for all callers. We propose that an ensemble classifier using the scores of multiple DNM callers would likely provide significant performance improvements, especially given the low concordance among individual callers.

The performance of the DNM callers is dependent on both the sequencing coverage and the VAFs of the DNMs. We observed clear distinctions in the patterns of VAF and read depth between true positive and false positive *dn*SNVs called by each caller. As noted earlier, both TrioDeNovo and DeNovoGear call large numbers of false positives. A considerable proportion of these calls are made within a very low VAF range and at a very high or low read depth, indicating technical artifacts such as sequencing errors. This is corroborated by investigating the trio VAF density patterns, which clearly show parental VAFs at or near 0 values for true positive *dn*SNVs, as they are absent in parents. In contrast, parental VAFs of false positive *dn*SNVs showed a large proportion at detectable VAF values. An exception to this was DeNovoCNN, which showed parental VAF levels similar to those seen in true positive dnSNVs - very low or zero - while the offspring VAFs were skewed towards very high values, up to 1, indicating potential homozygosity. Investigation of these variants indicated potential errors due to low read quality.

The VAF and read depth patterns of false positive *dn*SNVs obtained by consensus of all five callers in the real 1000G CEU dataset were similar to those of the true positive *dn*SNVs. It is possible that some of the false DNMs that were called by all the callers could be real somatic DNMs that were not validated by the original DNM study^32^. This is a limitation of our study that relies on this experimentally validated set of DNMs.

Another limitation of our comparative analysis is that we did not incorporate phased genotype information, even though some tools (e.g., DeNovoGear and PhaseByTransmission) are capable of providing this. This decision was made to ensure consistency across all DNM callers as most do not output phased genotypes. As a result, we were unable to assess parent-of-origin or haplotype context, which can provide valuable insights into the inheritance patterns of DNMs. Incorporating phasing in future analyses may improve the interpretability of DNM calls, especially in complex cases such as recurrent mutations or mosaicism.

The use of confidence scores from the callers could be an appropriate quality control step to filter out false positive DNMs. For DeNovoGear (PPDM threshold) and VarScan 2 (p-value), there were clear distinctions in confidence scores between true positive and false positive DNMs. In the case of DeNovoGear, a PPDM threshold of 0.99 yielded the majority of true positives. For VarScan 2, all true positive calls have p-value < 9.4048 x 10^-4^. However, for TrioDeNovo, DeNovoCNN and PhaseByTransmission, the distinction in confidence scores between true and false positive DNMs is unclear.

Many research groups have already incorporated standard variant processing pipelines, such as GATK, as part of their routine WGS data analyses. Hence, they may have VCF output files readily available for further processing. In such situations, it may be beneficial to use VCF-based input DNM callers such as TrioDeNovo, DeNovoCNN and PhaseByTransmission. On the other hand, DeNovoGear and VarScan 2 have only one pre-processing step; that of generating mpileup files (BCFs) from BAMs using a samtools mpileup command, which is an additional step in the pipeline.

Recently, collaborative efforts such as the Genome in a Bottle (GIAB) consortium have aimed to establish a reference standard for human genome sequence data^38^. Data for this purpose has been collected using 12 different sequencing technologies and library preparation methods. While gold standard calls have been generated for the Ashkenazim Trio data, these consist of gold standard truth variants within specific, high-confidence regions of the GIAB genome. There has not been a focus on generating a set of gold standard DNMs across the entire genome. There is a strong need for such public datasets containing validated DNMs, which could be extremely beneficial for the evaluation of existing callers and pipelines, as well as for the development of new algorithms.

Finally, the presence of post-zygotic DNMs in developmental disorders has recently received attention^39^. Determining the proportion of affected cells and tissue types can help to determine the timing of formation of DNMs, which plays a crucial role in the clinical phenotype. In the future, researchers will need to develop computational methods that can distinguish between germline and post-zygotic DNMs. This will require high-depth sequencing and computational methods that can confidently analyse the distribution of variant allele frequencies to detect differences between germline and post-zygotic DNMs that occur early enough in development to reach elevated levels.

## CONCLUSION

In this comprehensive evaluation of five DNM callers applied to WGS trio data, we reveal considerable variability in overall performance. Despite high sensitivity, the tools display low concordance and high numbers of false positive calls, highlighting challenges in accurate DNM detection. Among the callers, DeNovoGear and DeNovoCNN demonstrated the highest F1 scores, although no tool achieved high performance, underscoring the need for further methodological improvements.

Our findings emphasise the critical role of optimising filtering thresholds as well as investigating quality metrics such as VAF and read depth. The observed differences in parental and offspring VAF distributions between true and false positives offer important clues for enhancing DNM calling models. Limitations in current benchmarking datasets, particularly the lack of comprehensive gold standard DNMs across the genome and exclusion of phased genotype information, constrain full assessment of caller performance.

In the future, ensemble approaches that integrate multiple caller outputs, alongside advanced methods to distinguish germline from post-zygotic mutations, hold promise for improving DNM detection accuracy. The development and public availability of high-confidence, genome-wide validated DNM datasets will be critical to facilitate method development and benchmarking.

Ultimately, improving DNM calling accuracy will strengthen the utility of WGS trio analyses in both research and clinical settings, enabling more reliable identification of pathogenic mutations and advancing our understanding of genetic disease.

## KEY POINTS

- This study systematically evaluated five DNM calling tools (DeNovoGear, VarScan 2, TrioDeNovo, PhaseByTransmission and DeNovoCNN) using real and simulated WGS trio data, highlighting low concordance among the tools and significant issues with false positives.
- DeNovoGear and DeNovoCNN were identified as the best performing DNM callers, though all tools had relatively low F1 scores. DeNovoGear achieved the highest F1 score on the real 1000G dataset, while DeNovoCNN had the highest F1 score on the simulated data.
- False positive calls were a major challenge, with the study suggesting that an ensemble classifier using multiple callers could improve performance.
- The study emphasises the importance of incorporating confidence scores from callers and recommends using them to filter false positives.
- This evaluation provides valuable recommendations for selecting DNM calling tools and highlights the need for standardised public datasets of validated DNMs for future tool benchmarking.

## DATA AVAILABILITY

The 1000 Genome CEU trio we used was downloaded from ftp://ftp.1000genomes.ebi.ac.uk/vol1/ftp/data_collections/illumina_platinum_pedigree/data/CEU/NA12878/

The validated DNMs were taken from Conrad *et al.*^32^. Our trio simulated framework is available at https://github.com/VCCRI/TrioSim.

## Supporting information

Supplementary Matereial

