## Supplementary Matereial for "Systematic evaluation of *de novo* mutation calling tools using whole genome sequencing data"

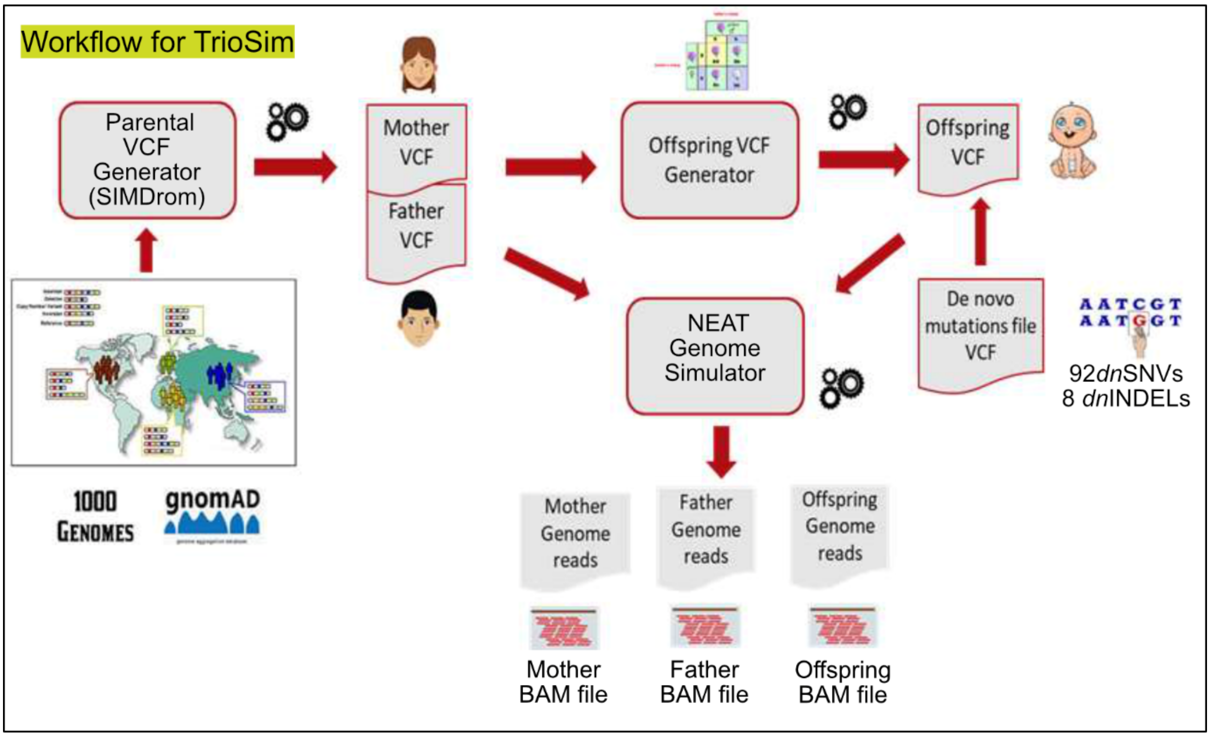


**Supplementary Figure S1. TrioSim - NEAT simulation WGS trio data.** Using an in-house application called "TrioSim", we created a synthetic WGS-based family trio set of BAMs. TrioSim utilised SIMdrom[33] to generate mother and father VCFs with allele frequencies observed in the European population data available in the 1000G project. The offspring VCF file was then generated by applying the Mendelian inheritance law to the merged parental VCF file, to infer the genotypes of the offspring. A set of 100 DNMS was spiked into the offspring VCF file to provide true-positive variants. Illumina-based NGS reads were simulated using the NEAT[35] open-source software package with the GRCh38 reference genome, at 30x coverage depth to produce three independent BAM files representing the trio (mother, father, and offspring). This all occurs within our in-house application TrioSim (at https://github.com/VCCRI/TrioSim).

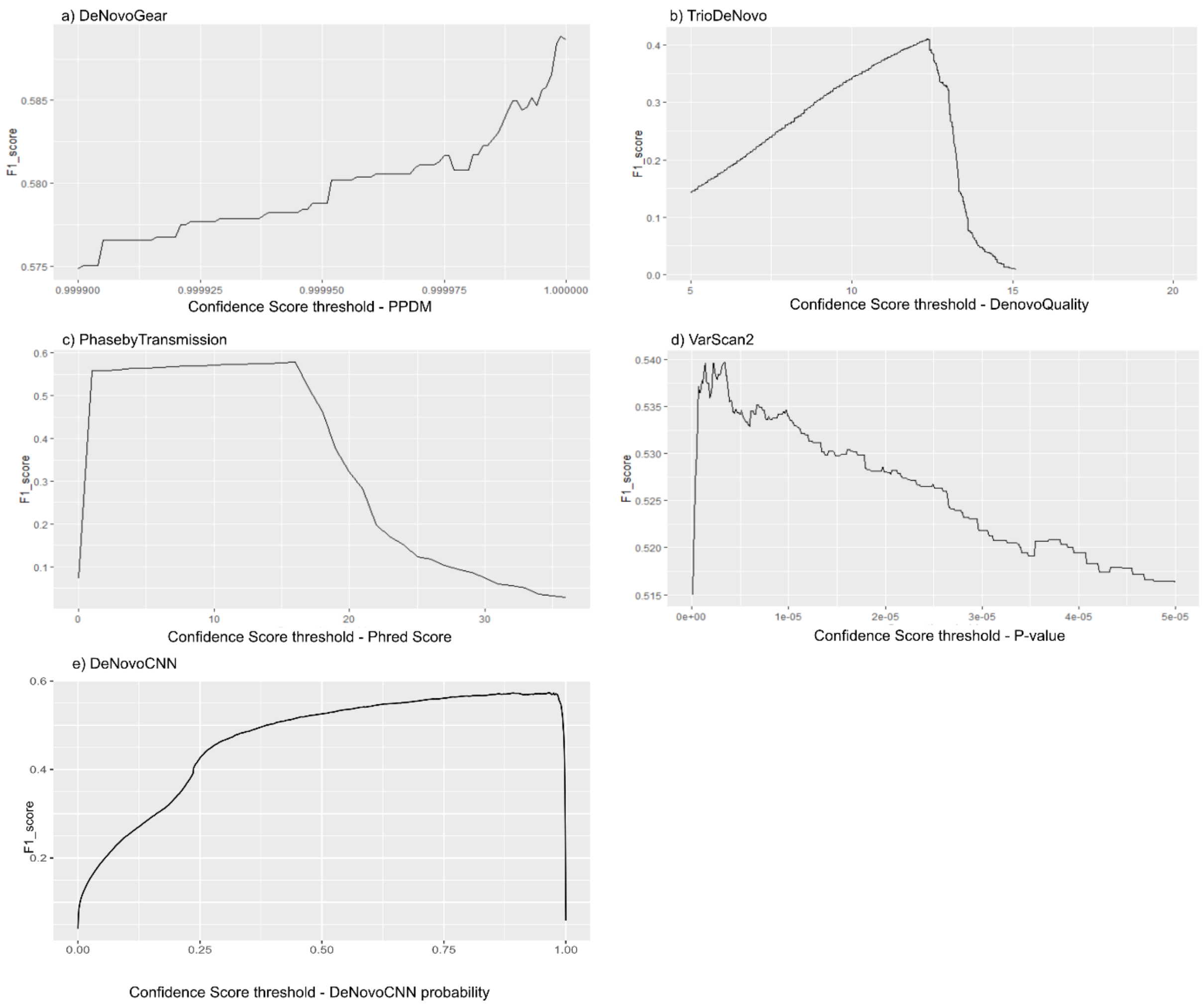


**Supplementary Figure S2. F1 scores for different confidence score threshold values for each DNM caller for the 1000g CEU trio dataset.** "Benchmarking-set maximal F1 thresholds" for DNM callers on 1000G CEU trio dataset (Table 5) were identified as the threshold resulting in the highest F1 score.

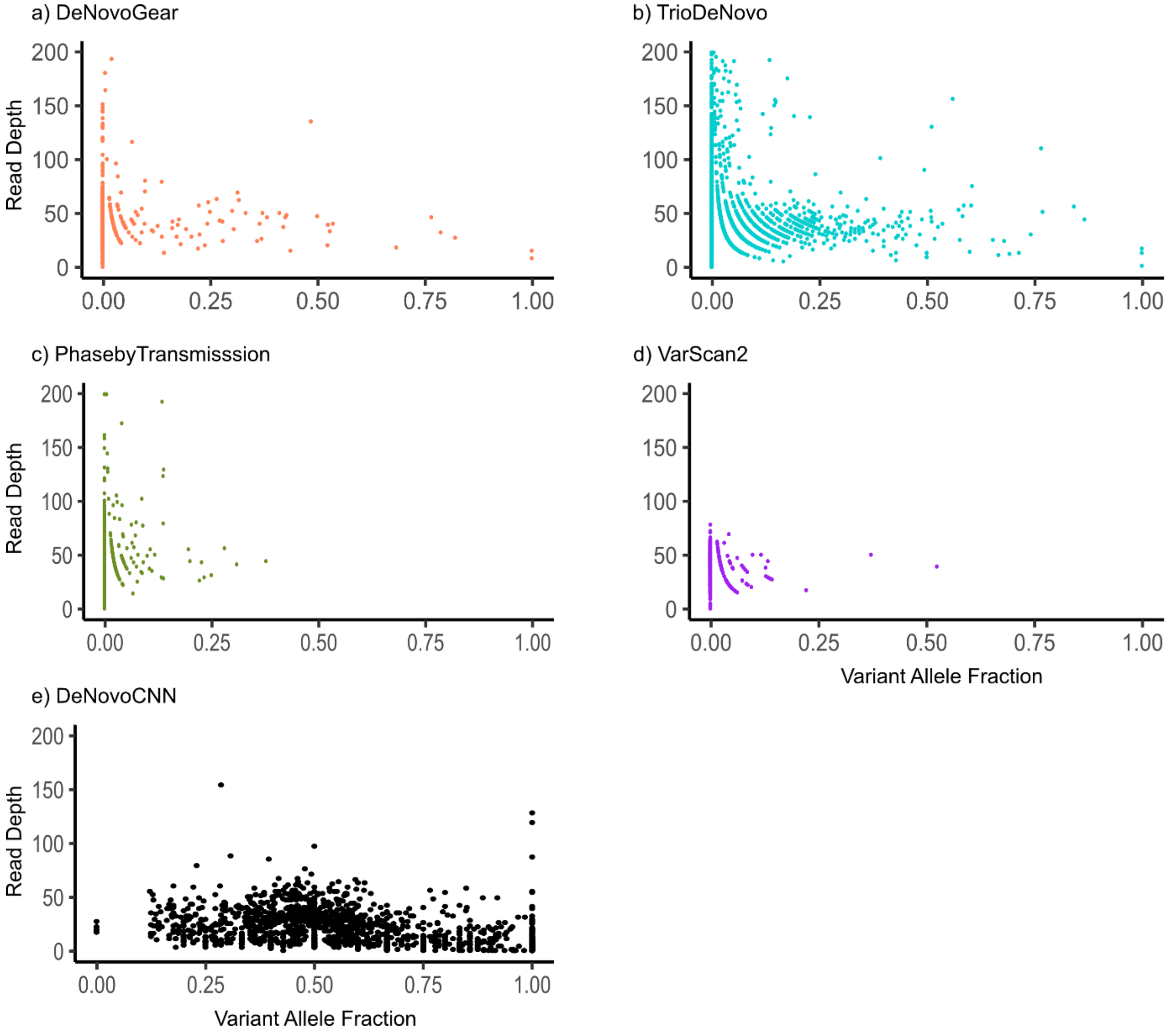


**Supplementary Figure S3. Variant Allele Frequency (or Fraction) vs. Read Depth for false positive dnINDELs for different DNM callers using CEU trio from 1000G.**

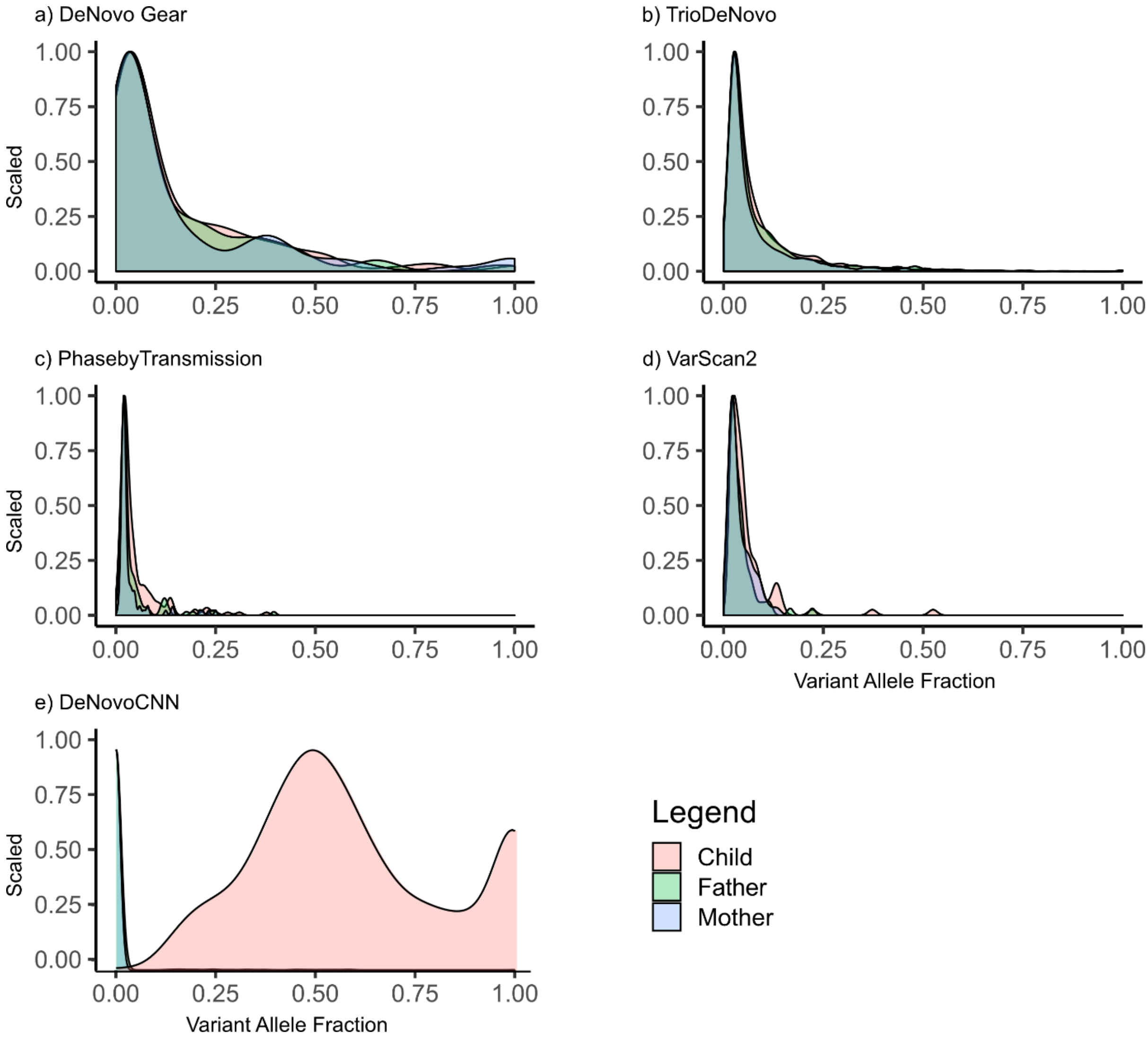


**Supplementary Figure S4. Density of VAF for dnINDELs for different DNM callers for CEU trio of 1000G.**

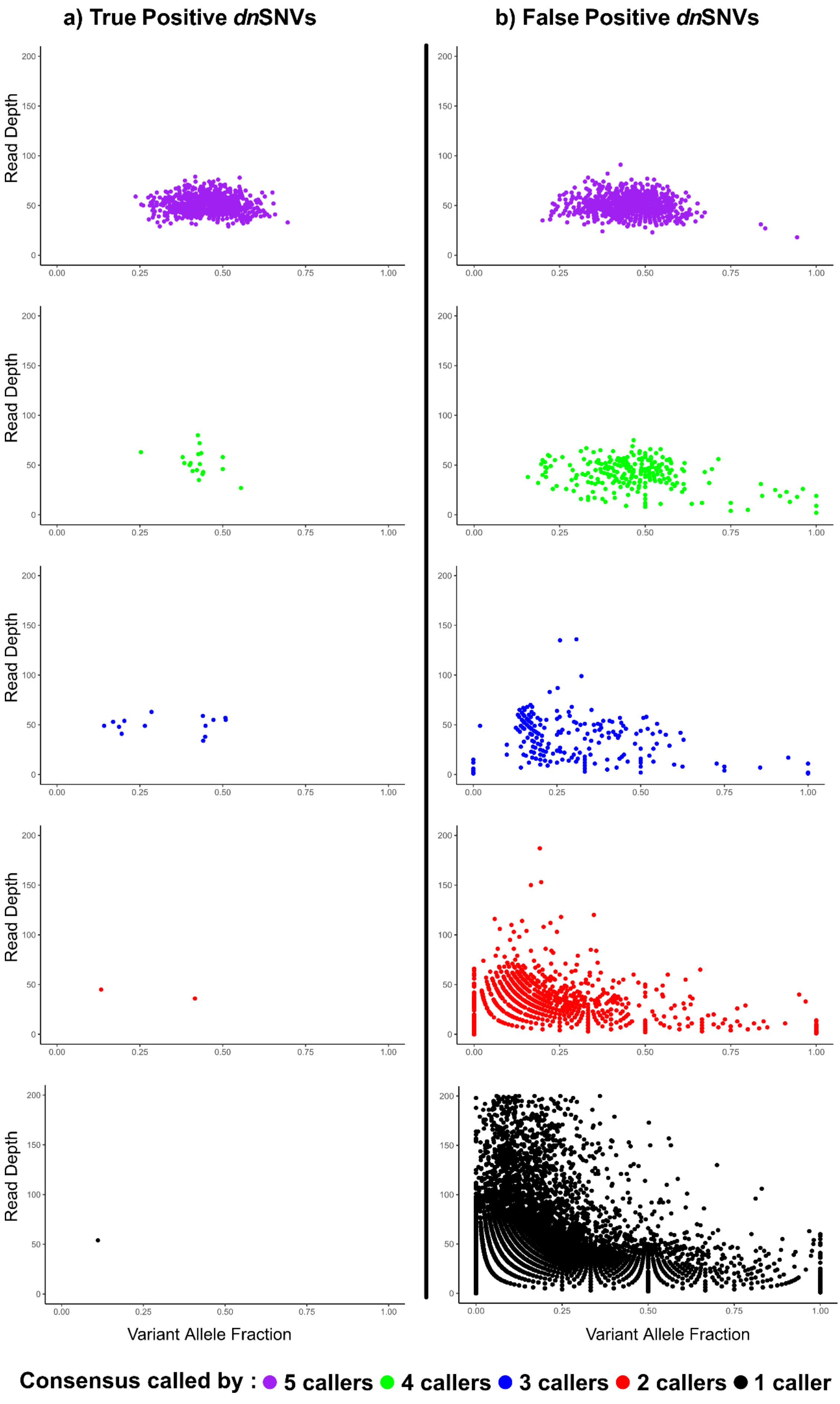

a) True Positive dnSNVs
b) False Positive dnSNVs
Read Depth
Variant Allele Fraction
0
50
100
150
200
0.00
0.25
0.50
0.75
1.00
Consensus called by : 5 callers 4 callers 3 callers 2 callers 1 caller

**Supplementary Figure S5**. **Variant Allele Frequency (or Fraction) vs. Read Depth for dnSNVs using the consensus of different numbers of DNM callers in CEU trio from 1000G.** True positive (a) and False positive (b) dnSNVs are shown. Different numbers of DNM callers are indicated by different colours.
